# Statistically inferred neuronal connections in subsampled neural networks strongly correlate with spike train covariance

**DOI:** 10.1101/2023.02.01.526673

**Authors:** Tong Liang, Braden A. W. Brinkman

## Abstract

Statistically inferred neuronal connections from observed spike train data are often skewed from ground truth by factors such as model mismatch, unobserved neurons, and limited data. Spike train covariances, sometimes referred to as “functional connections,” are often used as a proxy for the connections between pairs of neurons, but reflect statistical relationships between neurons, not anatomical connections, and moreover are not casual. Connections inferred by maximum likelihood inference, by contrast, can be constrained to be causal. However, we show in this work that the inferred connections in spontaneously active networks modeled by stochastic leaky integrate-and-fire networks strongly reflect covariances between neurons, not causal information, when many neurons are unobserved or when neurons are weakly coupled. This phenomenon occurs across different network structures, including random networks and balanced excitatory-inhibitory networks.

---

Identifying the strength and timescales of synaptic transmission between neuron pairs is a major goal of neuroscience, as it would greatly facilitate our understanding of how a neural circuit’s computational properties are shaped by its structure. It is now possible to record simultaneous activity from large populations of neurons, enabling the use of statistical methods to infer interactions between neuron pairs [1–4]. This has become a foundational tool for understanding the encoding and decoding properties of many biological neural networks. However, because no *in vivo* recording technique can record from all neurons in a circuit (Fig. 1) the inferred connections between neurons may only reflect statistical relationships between neurons, influenced by, but not necessarily representative of, the underlying anatomical connections [5–8].

**FIG. 1.**
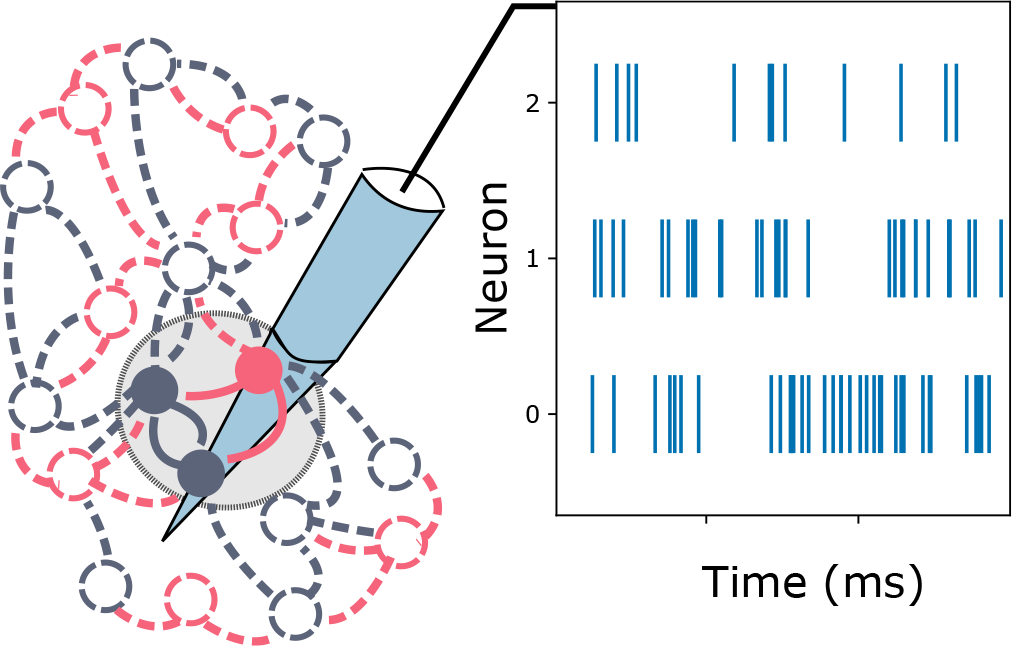
Schematics of the hidden neuron problem and effective neuronal connection inference. Recording techniques are often only able to record from a subset of neurons in a neural circuit, especially in the tissue of a living organism. In this schematic three neurons’ spike trains are shown to be recorded, while the activity of other nearby neurons remains unobserved. Statistical inference techniques applied to this activity data can predict effective connections between neurons, but these do not necessarily reflect the true anatomical connections.

To this end, it is useful to distinguish between two prominent measures of “functional” or “effective” interactions between neurons. “Functional” connections between neurons are estimated using pairwise cross-covariances between neuron spike trains, or other measures of neural activity, such as BOLD signals in fMRI [9]. Cross-covariances are not causal functions, and extracting information about causal circuit responses from them is not always possible. In contrast, “effective” interactions obtained by performing, e.g., maximum likelihood inference on neural activity data, can be constrained to be causal functions, and could therefore represent actual causal responses of neurons to spikes from pre-synaptic partners. Accordingly, effective interactions should be more useful for understanding how the underlying dynamics of neurons implement computations. However, in this work we use a combination of simulations and analytically tractable cases to show that when only a few neurons are recorded in a neural circuit — the typical case in any *in vivo* recording — the effective filters inferred from spontaneous neural activity correlate strongly with the causal half of the underlying spike train covariances.

## Results

Our goal is to understand how the circuit properties inferred using maximum likelihood inference are related to the ground truth properties of the network or the statistics of spontaneous neural activity—that is, in this work we do not consider stimulus-driven activity. We require both a generative model and an inference model. We choose to use the same model family for both, a generalized linear point process model (GLM). As a generative model the GLM can be interpreted as a leaky integrate-and-fire model in which spike emission is stochastic [10–13]. As an inference model the GLM has been used to fit neural spiking data from many brain areas, including the retina [1] and the lateral intraparietal area of macaques [14].

The GLM models spike train emission as an inhomogeneous Poisson process in which the instantaneous firing rate is conditioned on the past history of neural activity in the network:

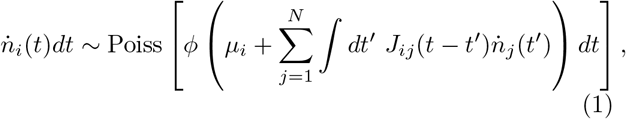

Where 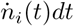 is the number of spikes neuron *i* fires within a small window [*t, t + dt*], ϕ (*x*) is a nonlinear activation function, *μ*_*i*_ is the baseline drive for neuron *i* that sets the baseline firing rate, and *J*_*ij*_(*t*) is the synaptic interaction or coupling filter from neuron *j* to neuron *i*. The parameters to be inferred from data are the baselines *μ*_*i*_ and synaptic filters *J*_*ij*_(*t−t*^*′*^); the nonlinearity *ϕ* (*x*) is often fixed, the canonical choice being an exponential, *ϕ* (*x*) ∝ exp(*x*) [1]. We will adopt this choice for our inference model, as it offers several simplifications in both our statistical inference procedure and mathematical analysis of the maximum likelihood procedure (but see the Supplementary Information [15] for a brief discussion of the expected effects of nonlinearity mismatch.). It is important to stress that when the number of observed neurons *N*_obs_ is not equal to the number of neurons N in the generative model we are dealing with a model mismatch problem and we do not expect statistical inference to recover the parameters of the generative model [13, 16]. To obtain the true generative model of just the observed neurons one should in principle marginalize out the unobserved neurons. This was done approximately by Ref. [13], but the resulting model is much more complex than the fully observed network model, and may not be suitable as a statistical inference model. It is therefore worth understanding how the inferred filters relate to the ground truth circuit properties when Eq. (1) is used as the inference model with *N*_obs_ < *N*.

The maximum likelihood estimates (MLE) of circuit properties are the model parameters that render the observed data as probable as possible under the inference model. These parameters can be found by maximizing the (log)-likelihood. For the GLM the log-likelihood is

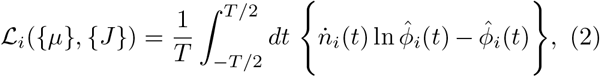

where *T* is the duration of the spike trains used in the fit and 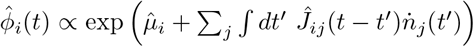. We use hats to distinguish parameters of the inference model from their corresponding ground truth counterparts. The parameters 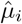 and 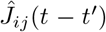 may be inferred independently for each neuron *i*.

We investigate the inferred synaptic interaction filters in two ways: i) performing maximum likelihood estimation on simulated data, mimicking analysis of real data, and ii) deriving an approximate system of equations for the inferred filters that we can study analytically. In both cases we focus on how the inferred filters change as a function of the number of neurons recorded.

### Simulations

We simulate two types of circuit networks in this first study. The first network type comprises random Gaussian networks in which*J*_*ij*_ (*t*) = 𝒥_*ij*_ *te*^*−t*/τ^ /τ ^2^ Θ(*t*), where the synaptic weights 𝒥_*ij*_ are normally distributed with zero mean and variance 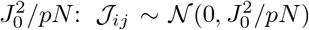. We set the baseline *μ*_*i*_ = −2 for all neurons and set the network sparsity *p* = 50%; for an exponential nonlinearity this allows us to tune the weight matrix coefficient *J*_*0*_ up to∼ 3.0 before the network becomes unstable. We note that unlike previous work that has analyzed the hidden neuron problem, we do not need to restrict our simulations or analysis to the weak-coupling regime [5].

Though random networks are commonly used in theoretical studies, they lacks some biological realism because every neuron can make synaptic both excitatory (positive) and inhibitory (negative) connections. We therefore also consider balanced networks of excitatory and inhibitory (E-I) populations, for which each neuron is either excitatory or inhibitory and only makes connections of a single sign [17, 18].

For each network type we simulate the spiking activity of 64 neurons for 2 million time points. While simulating larger networks is possible, fitting to larger networks is very memory intensive. To fit the GLM to these simulated data sets it is necessary to parameterize the filters *Ĵ*_*ij*_ (*t*), either by inferring the value of the filter at each time point (requiring as many parameters as the number time-bins used to represent the filter) or by representing them as weighted sums of basis functions and inferring the unknown weights. The basis function approach reduces the number of unknowns to the number of basis functions used, which requires less data than inferring each time point. The filter shapes that can be inferred are constrained by one’s choice of basis functions, whereas inferring each time-point can represent any function given enough temporal resolution and data, but are often noisy estimates without adding fit penalties to impose smoothness. In practice, the basis function representation is preferred, but for our analyses the time-point inference will reveal interesting relationships between the inferred filters and statistics of the circuit activity. For this reason we also do not impose any regularization on the basis-less fit to smooth out the inferred filters.

In Fig. 2 we show the results of statistical inference on 3 out of 64 neurons from the random Gaussian network. The results show that the inferred filters do not match the ground truth filters of the generative model (blue dashed line), regardless of whether we use basis functions to infer the filters (solid red lines) or infer each time point (red dots). However, we observe that the basis-less inferred filters do match quite well with the empirical cross-covariances of the spike train pairs (grey bars). Since the MLE inferred filters and spike train covariances are estimated using two independent methods, it is surprising to observe such a strong correlation between them.

**FIG. 2.**
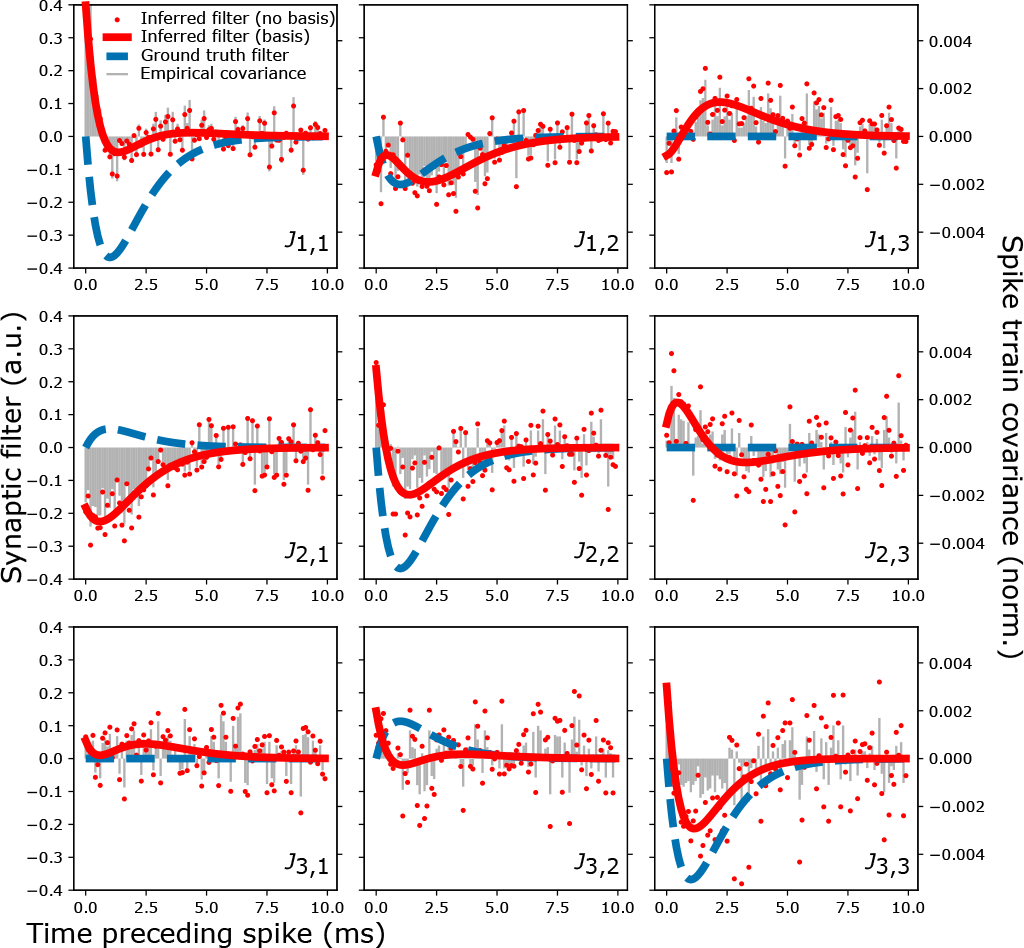
Maximum-Likelihood estimates (MLE) of cou pling filters *J*_*ij*_ for 3 neurons out of 64 in a circuit with 50% sparsity and Gaussian weights. The ground truth synaptic connection filters of the generative model are shown in blue dashed lines (values on left axis). Two types of inferred filters are shown: for each time point (red points) and using a basis expansion (red solid line). We also compare the filters to the empirical covariances, shown in grey bars (values on right axis). The covariances correlate strongly with the filters inferred without using a basis expansion.

To verify the correlation is robust and not just a visual artifact, we calculate the linear Pearson correlation coefficient between inferred filters and the empirical spike-train covariances, varying the number of observed neurons in both random and E-I networks. The distribution of the correlation coefficients as a function of the fraction of neurons observed is shown in Fig. 3A for the sparse random networks. On the left panel, we see that the correlation between auto-covariances and self-coupling filters is generally very close to 1 when. ≲10% of the network is observed, dropping to a median of ∼0.7 when the network is fully observed. On the right panel, the cross-covariances between different neurons are all even more strongly correlated with their corresponding inferred cross-coupling filters. Same trends are shown for the balanced E-I networks in Fig. 3B.

**FIG. 3.**
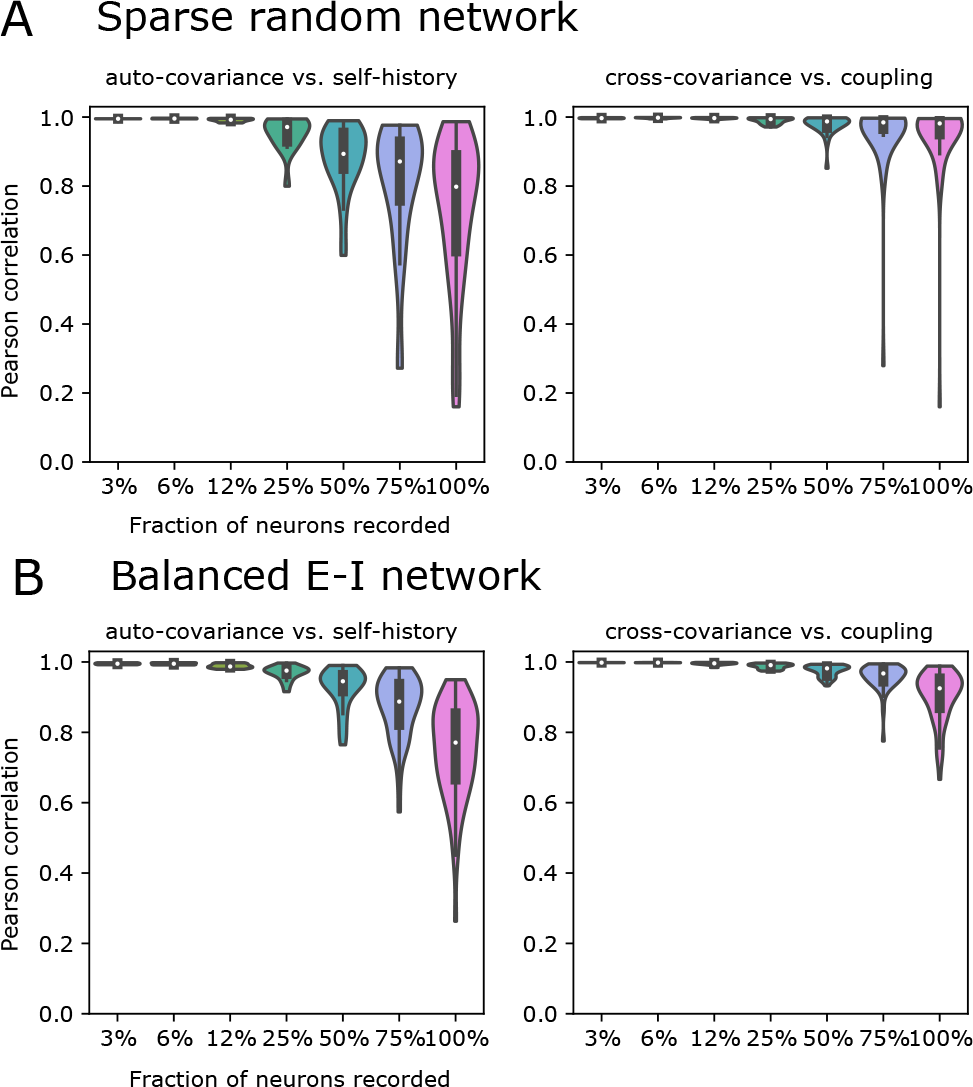
Pearson correlation between the spike train correlation and MLE inferred coupling filters in strongly coupled random networks and E-I balanced networks of 64 neurons. **A**. Violin plots show how the correlations change when the different number of neurons are observed and different amount of spike train data is used in inference. 3%, 6%, 12%, 25%, 50%, 75%, and 100% percentage of observed neurons in this 64 neuron network corresponds to 2, 4, 8, 16, 32, 48, and 64 neurons being observed. The spike train correlation functions strongly correlate with the MLE inferred filters inferred for each time point when less neurons are observed and the correlation decreases as more neurons are observed. The **left panel** shows the distribution of the Pearson correlation coefficients between the autocovariance and self-history coupling filters, while the **right panel** shows the distribution of the Pearson correlation coefficients between the cross-covariance and coupling filters between neuron pairs. **B**. Same as **A** but for the balanced E-I networks.

We also investigated how the strength of these correlations depends on the synaptic weights. For random Gaussian networks we varied the standard deviation of the weights, *J*_*0*∈_*{*1, 2, 3*}*, while for balanced E-I networks we varied both excitatory and inhibitory weights by a multiplicative factor *J*_0_ ∈ *{*1, 4, 7*}*. The results shown in Fig. 3 correspond to the strongest synaptic strengths we investigated, namely, *J*_0_ = 3 for the random networks and *J*_*0*_ = 7 for the E-I networks. We find that correlations between covariances grow even stronger when the synaptic connections are weaker, shown in Fig. 4. As shown in the right panel of Fig. 4A, in the weak coupling limit the Pearson correlations between the MLE inferred filters and the spike train covariances are high even if all the neurons in the network are observed. Smaller *J*_0_ confined the network to a noise-driven regime, where each neuron’s firing rate is dominated by the same baseline drive *μ* set in the generative model, and a higher *J*_0_ tunes the network into a strong coupling regime with more variable firing rates across neurons.

**FIG. 4.**
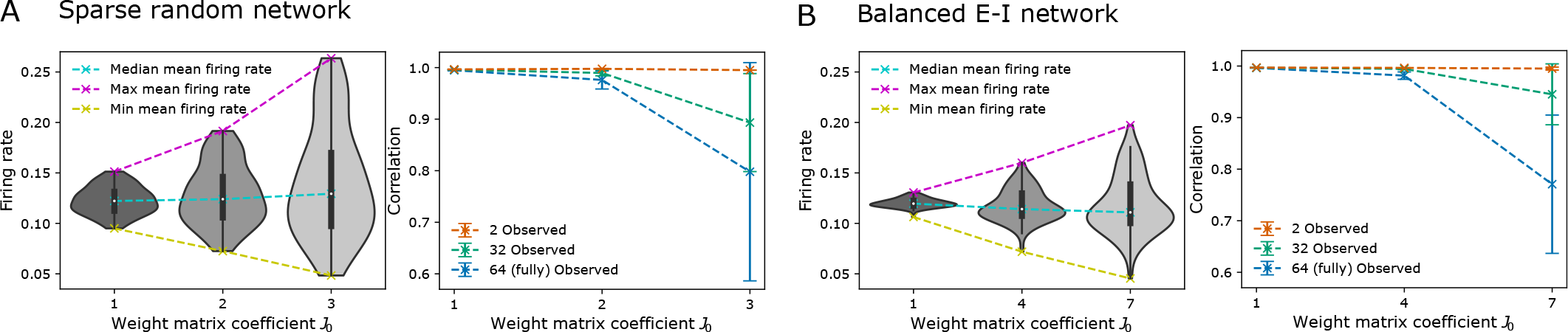
Correlations are high for both random and balanced excitatory-inhibitory networks in weak coupling regimes. **A**. Random network. When the weight matrix coefficient *J*_0_ decreased from 3 to 1, the network transitioned into a noise-driven regime, and the mean firing rates of the neurons varied less and were driven by the same baseline drive in the generative model, as shown in the **left panel**. As shown in the **right panel**, in the weak coupling regime, such as when *J*_0_ = 1, high correlations between the MLE inferred filters and spike train covariances were observed even when all the neurons in the network are observed, as compared to the network in a strong coupling regime where the high correlation between the MLE inferred filters and spike train covariances only happen in the sub-sampled network. **B**. Similar results hold for a balanced excitatory-inhibitory network, where 20% of the neurons are inhibitory and 80% of them are excitatory. The weight matrix coefficient *J*_0_ is tuned from 1 to 7, with *J*_0_ = 7 the largest possible integer value for which the spiking process is still stable.

Because our simulation results demonstrate a high degree of correlation between the inferred synaptic filters (without using basis functions) and the empirically estimated spike-train covariances, some property of the network statistics or maximum likelihood inference procedure must give rise to these strong correlations when the network is subsampled. To better understand this relationship, we turn to an analytic analysis of the maximum likelihood estimation of subsampled networks.

### Analytical analysis

We will first derive the MLE equations for the spiking network model with an exponential nonlinearity, and then approximate the spike trains as a Gaussian process to analytically solve the MLE equations for some simple networks, which elucidates how the inferred filters are related to the spike train covariances. We can study the maximum likelihood equations directly in continuous time. In the limit of infinitely long spike trains (*T*→), the right hand side of Eq. (2) becomes an expectation over the ground truth spike train process for circuits in which the spontaneous activity settles to a steady state.

The maximum likelihood equations are obtained by taking derivatives of the log-likelihood with respect to 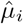 and *Ĵ*_*ij*_ (*t)*, the parameters to be inferred. For the exponential nonlinearity this gives a set of self-consistent equations to be solved,

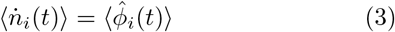

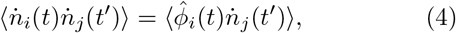

where the angled brackets denote an expectation over spike trains of the *generative* circuit model, not the inferrence model. In a stationary steady-state Eq. 3 will be independent of time t and Eq. 4 will depend only on the time difference *t −t*^*′*^.

We exploit the choice of the exponential nonlinearity to relate the expectations in Eqs. (3)-(4) to the moment generating functional of the spike train process:

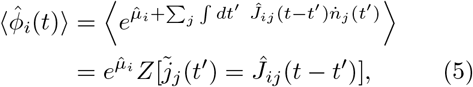

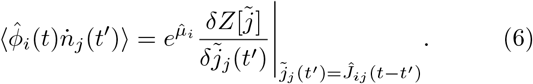

where *Z* is the moment generating functional of the fluctuations, defined for an arbitrary “source” variable 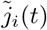 as 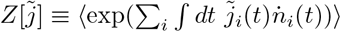.

An important implication of Eqs. (5)-(6) is that the moment generating functional contains only information about the non-causal statistical *moments* of the spike train process; they do not directly contain any information about *response functions*, which are causal. In a path integral formulation of this stochastic process, one can formulate a more general moment generating functional that contains information about both the statistical moments and the causal response functions of the process. Crucially, however, the information about the response functions drops out of the expectations we have computed, meaning that the MLE equations do not directly contain any information about the response functions. In fully observed systems in steady-states it is often possible to derive fluctuation-dissipation relationships between the statistical moments and response functions [19]. While our results suggest that such a relationship may enable the accurate inferrence of the ground truth connections in a fully observed circuit, inference from subsets of neurons cannot recover information about circuit response functions, and the inferred connections may reflect only covariances in neural activity.

The moment-generating process *Z* for the spiking network cannot be solved in closed form, so to glean some concrete insight about the inferred circuit parameters from Eqs. 3 and 4 we approximate the fluctuations *δn*_*i*_*(t)* as a Gaussian process, which amounts to neglecting the contribution of all statistical moments beyond pairwise statistics. We then use Eq. 5 to eliminate the dependence on 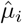 in Eq. 6, leaving a system of Wiener-Hopf integral equations to solve for the filters *Ĵ*_*ij*_ (*t)*:

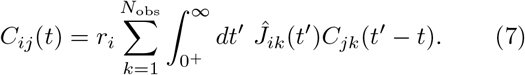

Here, 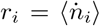 is the mean firing rate of neuron *i* and 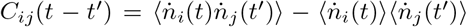 is the spiketrain covariance. In practice these quantities would be estimated from data and hence known, but in this analysis we will estimate them using a mean-field analysis of the spiking network model [12, 13].

Although Eq. 7 looks like it can be solved using a Fourier transform, it is only valid for *t* > 0. If this restriction is neglected, the solutions *Ĵ*_*ij*_ (*t)* may be non-causal. Solving this system of equations while imposing causality is difficult and an area of active research [20]. To study a tractable case, we consider a homogeneous network of all-to-all coupled neurons, which displays the key features observed in our simulations. We choose *J*_*ij*_*(t) = Jt exp(− t/τ)Θ (t)/ τ* ^*2*^, including the self couplings *i* = *j*, and homogeneous baselines *μ*_*i*_ *= μ*. The self-consistently calculated mean firing rates can be solved in terms of the Lambert W function,*r* = *−(N J)* ^*−1*^ *W* _*−1*_ *(−N J λ* _*0*_ *e* ^*μ*^), defined as the solution of the transcendental equation *x* = *W*_*−*1_(*x*) exp(*W*_*−*1_(*x*)) for which 1/*e* < *x* < 0. This restriction defines the branch of the Lambert W function that we must use for excitatory *J* > 0. It follows that the network is only stable for *N J λ*_*0*_ exp(1 + *μ*)) < 1.

The mean-field (or more precisely, “tree-level”) approximation of the spike train covariance is

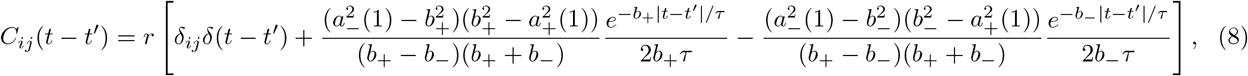

where 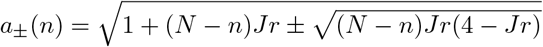 and 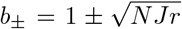. With this result, we can use the Wiener-Hopf procedure [20] to solve Eq. for the selfhistory filter of *N*_obs_ observed neurons. We find

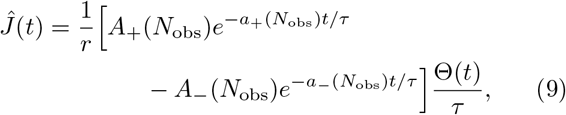

where *A*_*±*_(*N* _obs_) depend on *a*_*±*_(*N*_obs_), *a*_*±*_(1), and *b*_*±*_. Their full expressions are given in the Supplementary Material, but for *N*_obs_ = 1 they reduce to 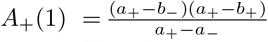 and 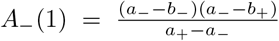. In the limit *N*_obs →_ *N* this filter recovers the ground-truth filter *J(t) = Jt exp*(− *t*/τ)Θ(*t*)/ *τ* ^2^. The inferred filter differs from the effective filters of the model in which all but the recorded neuron are marginalized over [13], due to the fact that our inference model does not include an effective Gaussian noise generated by the fluctuations of the unobserved neurons’ activity. Models with common Gaussian driving noise have been fit to data [21], but the fitting procedure is more involved than standard maximum likelihood inference, and we do not consider it here. In Fig. 5A, we plot the filters for different numbers of observed neurons *N*_obs_, observing that the amplitude and decay rate of the filters decreases as *N*_obs_ increases, for both fits to simulated data and our solutions of Eq. (7) (inset). The mean-field theory correctly captures the qualitative trend. In the homogeneous network the self-history filters *Ĵ* _*i i*_ (*t*) = *Ĵ* _self_ (*t*) and crosscoupling filters *Ĵ* _*i*≠*j*_ (*t*) = *Ĵ* _cross_ (*t*)are identical, though the inferred filters differ slightly due to finite data effects.

**FIG. 5.**
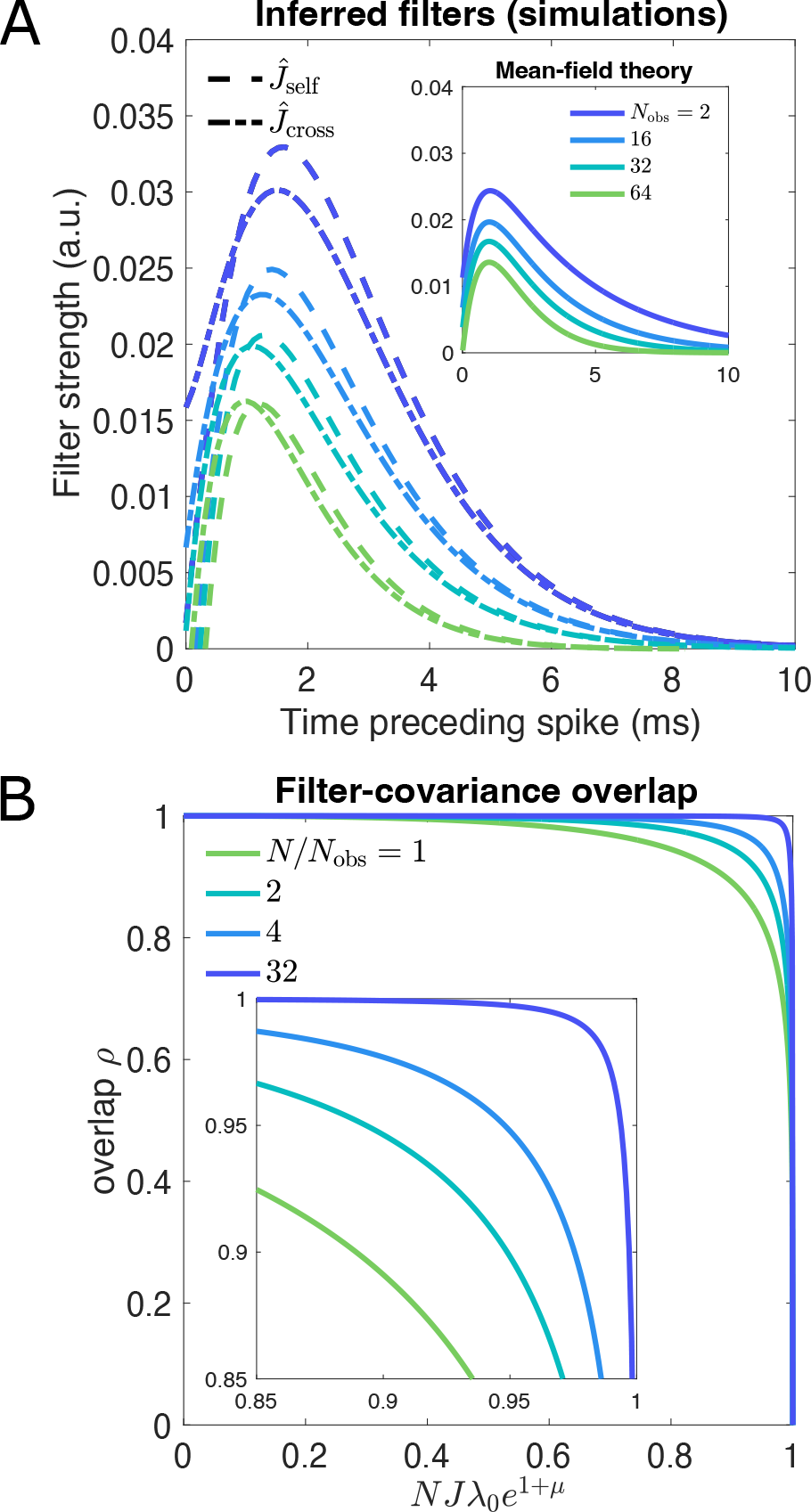
Dependence of fits on the number of observed neurons in a homogeneous network. **A**. The inferred filters decrease in amplitude as the number of observed neurons *N*_obs_ increases to the total number of neurons in the network *N*, observed in fits to simulated data and our mean-field analysis (inset). In the ground truth model the self-history filters *Ĵ* _self_ (*t*) and cross-coupling filters *Ĵ* _cross_ (*t*) are identical, as is the theoretical prediction from solving Eq. (7). The inferred selfand cross-coupling filters differ due to finite data effects, but are close. Simulation parameters: *N* = 64, *J* = 0.037, *μ* = *−*2, *λ*_0_ = 1, 4 10^6^ time points. **B**. The predicted overlap drops as *N J λ* _*0*_*e*^*1+μ*^ increases, with varying speed depending on the ratio *N/N*_obs_. *N/N*_obs_ = 1 corresponds to the fully observed case. The inset shows an enlarged view of the drop. Compare to Fig. 3.

We quantify the overlap coefficient between the inferred filter *Ĵ* (*t*) and the auto-covariance *C(t)*:

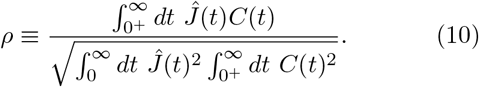

The overlap is equivalent to the Pearson correlation coefficient estimated from infinite time-points. The general expression is rather unwieldy, but it can be expressed as a function of the combined parameter *N J λ* _*0*_ *e*^*1+μ*^ ≤ 1, where 1 is the edge of stability of the network. As shown in Fig. 5B for various values of *N*/*N*_obs_, we observe that the overlap is generally very high away from the edge of stability of the network, but drops to 0 as the edge of stability is approached, capturing the behavior observed in Fig. 3. For large networks and a small fraction of recorded neurons (*N » N*_obs_) close to the edge of stability of the network (*N J λ* _*0*_ *e*^*1+μ*^ → 1 *−)* the overlap is asymptotically

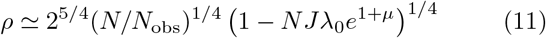

The window 1 *−N J λ* _*0*_ *e*^*1+μ*^ over which the correlation between *Ĵ* (*t*) and *C(t)* drops from *ρ* = 1 to 0 is *𝒪* (*N*_obs_/*N*). Thus, even when the synaptic strength is quite strong, a heavily subsampled network must be tuned extremely close to the edge of stability before the inferred filters and spike train covariances differ appreciably.

### Discussion and future directions

The homogeneous network yields valuable insight into what is occurring in our simulations of random networks and balanced excitatory-inhibitory networks. The inferred filters obtained by maximum likelihood estimation are shaped by network responses only through the non-causal spike-train covariances. When the network is subsampled it may not be possible to recover the causal response functions from activity covariance, and imposing causality on the inferred filters does not directly reflect the causality of neural responses. This suggests that the synaptic connections inferred from spontaneous activity data may not offer any advantages over the cross-covariances (“functional connections”).

Mapping out the network structure and inferring connections between neuron pairs from the recorded spike train data are challenging tasks, and understanding the results one obtains requires careful consideration of the assumptions underlying the statistical models fit to data. For example, Refs. [22] and [23] showed that it is possible to reconstruct neuronal connections from spike train covariances in certain sparse networks. While some efforts have been made to infer neuronal connections between observed and hidden neurons, these methods must often make unrealistic assumptions, like allowing allow acausal connections between the observed and the hidden neurons [24], the number of hidden neurons is less than the observed neurons [25, 26], or require careful modeling of the hidden neuron populations [27].

Other data-driven methods have emerged for inferring putative causal flows of information in neural circuitry, such as Granger causality, information-theoretic measures, or novel sampling paradigms [2, 4, 7, 28–32]. However, even causality tests like Granger causality may not identify true causal influences between neurons due to unobserved neurons [33]. Theoretical and simulation-based analyses like Refs. [5], [16], and this work are needed to understand the limits of statistical inference on subsampled neural data.

In this work we focused on analyzing maximum likelihood inference applied to spontaneous activity data. However, applications of this method to real data often involve stimulus-driven activity [8]. One might wonder whether such input drives would result in maximum likelihood estimates that capture true causal activity within a circuit. The answer is no if the stimulus is provided as an input over a long single trial, or if it is chosen randomly across multiple trials, as in that case the stimulus can be treated as an additional stochastic process, and we expect the maximum likelihood estimates to reflect stimulus-spike covariances, not response functions. However, if an intra-cortical perturbation is delivered to a circuit and repeated several times after the activity returns to its steady state, then it becomes possible to align data to these events and estimate network response functions. These perturbations can be explicitly included in the likelihood of the model and may enable inference of synaptic filters *Ĵ* _*ij*_ (*t*) that reflect true causal flows of information through the network. Responses to perturbations have been gaining traction in neuroscience [34–36], and extensions of the analyses presented here will be valuable in guiding this next phase of probing neural circuitry.

## Acknowledgments

We thank Il Memming Park, Yuan Zhao, and Matthew Dowling for helpful discussions, and Siddharth Paliwal for code to generate the E-I networks.

