## Supplementary information for "Statistically inferred neuronal connections in subsampled neural networks strongly correlate with spike train covariance"

### Appendix A: Methods

#### 1. Spike train simulation with a linear-nonlinear Poisson cascade model

In this work, we use a generalized linear model (GLM) to simulate the neuron spike trains. In the GLM generative model, neurons emit spikes probabilistically following a Poisson process, with the rate given by $\phi\left(\mu_i + \sum_j \int dt'\ J_{ij}(t-t')\dot{n}_j(t')\right)$. Discrete spikes are generated for a small observation window $dt = 0.1$. A first-order alpha function $J_{ij}(t) = \mathcal{J}_{ij}\ t\ e^{-t/\tau}/\tau^2\Theta(t)$ is used as the ground-truth interaction filters that govern the interaction of neuron $j$'s spike train history on neuron $i$'s instantaneous firing rate. Causality is imposed through the Heaviside step function $\Theta(t) = 1$ if $t > 0$ and 0 otherwise. The weight matrix $\mathbf{J}$ with entry $\mathcal{J}_{ij}$ sets the interaction strength of the filters and $\tau = 1$ sets the typical timescale of the decay of the response. The spike train is simulated by solving a second-order differential equation with a 4th-order Runge-Kutta method to conveniently track the spike train history, following the method used in previous works [12, 13, 37]. While simulating the spike train data, we run the simulation up to 2 million observation windows, and use different amounts of the spike train data in inferring the coupling filters, e.g. in Fig. 3.

*Random network weight matrix generation.* Following previous work [12, 13], we generated a 64-neuron random network weight matrix with a sparsity $p = 50\%$, so that only half of the connections are non-zero. The non-zero synaptic weight strengths were drawn independently from a normal distribution with zero mean and standard deviation $J_0/\sqrt{pN}$, where $J_0$ is the weight matrix coefficient and $N$ is the number of neurons in the network. The weight matrix coefficient $J_0$ takes three values 1, 2, 3, while we set the baseline drive $\mu_i = -2$ throughout the study. $J_0 = 3$ is the largest integer value we can set to still have a stable spike train process in the simulation, thus the network is considered to be in a high coupling regime in that case. Importantly, the diagonal entries of the weight matrix were always set to $-1$ for different $J_0$ to simulate a soft refractory period for the neurons from their own spike history.

*EI network weight matrix generation.* Following previous works [17, 18], we generated a 64-neuron excitatory-inhibitory (EI) network weight matrix with 20% (13) inhibitory neurons and 80% (51) excitatory neurons. Excitatory neurons make connections to excitatory neurons with a probability 20% and other all other neuron pairs (E-I and I-I) make connections with a probability 50%. The weight matrix coefficient $J_0$ was set to be a multiplicative factor of the weight matrix. The base weight of the excitatory connections was set to 0.04125 and the weight of the inhibitory connections was set to $-0.16625$ when $J_0 = 1$. Thus for the largest possible integer weight matrix coefficient $J_0 = 7$ in Fig. 4D-F, the excitatory and inhibitory weights were 0.28875 and $-1.16375$, respectively. The diagonal entries were always set to $-1$ for different $J_0$ to simulate a soft refractory period for the neurons from their own spike history.

#### 2. Neuronal connection inference with maximum likelihood estimation

We infer the neuronal connections based on a generalized linear model with an exponential inverse-link function, which amounts to a model-matched inference given the same model used in generating the spike trains. The observed neuron spike train $\{\dot{n}_i(t)\}$ are assumed to follow $\dot{n}_i(t)dt \sim \text{Poiss}[\phi(\hat{\mu}_i + \sum_{j\in obs} \int dt'\ \hat{J}_{ij}(t-t')\dot{n}_j(t'))dt]$, where $\hat{\mu}_i$ and $\hat{J}_{ij}$ are the inferred baseline drive and interaction filters to be determined and the observation window $dt$ is set to 0.1. The likelihood function is thus

$$L_i(\hat{\mu}, \hat{J}) = \text{Prob}(\{\dot{n}_i(t)\}|\hat{\mu}_i, \hat{J}_{ij}) = \prod_t \frac{(\phi_i(t)dt)^{\dot{n}_i(t)dt}}{(\dot{n}_i(t)dt)!} e^{-\phi_i(t)dt}. \quad \text{(A1)}$$

We use the Tweedie regressor with power 1 and log-link function in the scikit-learn (v.0.24.2) package to perform the inference [38], which minimizes the unit deviance and can be shown to be equivalent to maximizing the likelihood function in Eq. A1. No regularization penalty is added for all the inference procedures used in this work.

For the inference of the filters with basis functions as shown in Fig. 1C, we use basis functions of the form

$$\alpha_n(t) = t^n \exp(-t/\tau)\Theta(t)/\tau^n,$$

for $n = 0$, 1, and 2. In this scenario, the number of unknowns for inferring the filters decreases to 3, the same as the number of basis functions. The inferred neuronal connections are truncated at 100 observation windows, corresponding to 10 s for the chosen time window $dt = 0.1$ s. These basis functions are motivated by theoretical work that suggests the linear spike train filters in the true effective model for only the observed neurons will be a series of such functions [13].

For the inference of the filters without using basis functions, we use the same number of 100 observation windows, and thus 100 unknowns must be inferred to determine the coupling filters at each time point preceding the spikes.

#### 3. MLE solution from the path integral formalism of spike train process

Maximizing the likelihood function in Eq. A1 amounts to solve for the zero points of its derivatives with respect to the unknowns $J_{ij}$ and $\hat{\mu}_i$ in $\phi_i(t) = \lambda_0 \exp(\hat{\mu}_i + \sum_j \hat{J}_{ij}\dot{n}_j)$. For mathematical simplicity, we take the logarithm of the likelihood to get the log-likelihood function $\mathcal{L}_i = \log(L_i)$ and maximize the log-likelihood,

$$\frac{\partial \mathcal{L}_i}{\partial \hat{\mu}_i} = \lim_{T\to\infty} \frac{1}{T}\int_{-T/2}^{T/2} dt \, \left(\frac{\dot{n}_i(t)}{\phi_i(t)} - 1\right) \partial_{\hat{\mu}_i}\phi_i(t) = \lim_{T\to\infty} \frac{1}{T}\int_{-T/2}^{T/2} dt \, \left(\frac{\dot{n}_i(t)}{\phi_i(t)} - 1\right) \phi_i(t) = 0, \tag{A2}$$

where we note that $\partial_{\hat{\mu}_i}\phi_i(t) = \phi_i(t)$ for the choice of exponential nonlinearity. For a stationary system the time average will tend to the expected value due to ergodicity, and is equivalent to forming the log-likelihood using a large number of independent trials, which limit to the expected value for infinitely many trials. Thus, Eq. A2 can be simplified to

$$\langle \dot{n}_i(t) \rangle = \langle \phi_i(t) \rangle \,, \tag{A3}$$

which will be independent of the time $t$ for a stationary process, which we assume the steady state to be.

Similarly,

$$\frac{\delta \mathcal{L}_i}{\delta \hat{J}_{ij}(t)} = \lim_{T\to\infty} \frac{1}{T}\int_{-T/2}^{T/2} dt' \, \left(\frac{\dot{n}_i(t')}{\phi_i(t')} - 1\right) \partial_{\hat{J}_{ij}(t)}\phi_i(t') = \lim_{T\to\infty} \frac{1}{T}\int_{-T/2}^{T/2} dt' \, \left(\frac{\dot{n}_i(t')}{\phi_i(t')} - 1\right) \phi_i(t')\dot{n}_j(t'-\tau) = 0, \tag{A4}$$

where we note that $\partial_{\hat{J}_{ij}(t)}\phi_i(t') = \phi_i(t')\dot{n}_j(t'-t)$ for the choice of exponential nonlinearity and thus the equation can be reduced to Eq. 4.

As explained in the main text, for an exponential nonlinearity we can relate the expectations over $\phi_i(t)$ to the moment-generating functional of the spiking process and set $\lambda_0 = 1$,

$$\langle \phi_i(t) \rangle = e^{\hat{\mu}_i} Z[\tilde{j}_i(t') = \hat{J}_{ij}(t-t')],$$
$$\langle \phi_i(t)\dot{n}_i(t-\tau) \rangle = e^{\hat{\mu}_i} \left.\frac{\delta Z[\tilde{j}]}{\delta \tilde{j}_j(t')}\right|_{\tilde{j}_j(t')=\hat{J}_{ij}(t-\tau)},$$

(Eqs. 5 and 6 in the main text), where $Z[\tilde{j}] \equiv \langle \exp(\sum_i \int dt \; \tilde{j}_i(t)\dot{n}_i(t)) \rangle$. The moment generating functional cannot generally be solved in closed form, so to make use of these equations we will need an approximation. We will use mean-field theory with Gaussian fluctuation corrections to approximate the spike trains as a Gaussian process, for which the moment generating functional is known in closed form.

Our calculation of the moment generating functional of the spike train process makes use of the path-integral formalism based on the spiking network model [12, 13, 39, 40]. Following Ocker *et al.* [12], we introduce an auxiliary variable $\tilde{n}$, called the "response variable," then and the action of the spike train process under our GLM neuron model becomes

$$S[\tilde{n}, \dot{n}] = \sum_i \int dt \, \left[\tilde{n}_i(t)\dot{n}_i(t) - (e^{\tilde{n}_i(t)} - 1)\phi\left(\mu_i + \sum_j (J_{ij} * \dot{n}_j)(t)\right)\right], \tag{A5}$$

such that the joint probability distribution of the spike train and auxiliary variable follows,

$$\text{Prob}[\tilde{n}, \dot{n}] \propto e^{-S[\tilde{n},\dot{n}]}. \tag{A6}$$

Going forward, we make a change of variables $\dot{n}_i = r_i + \delta n_i$ where $r_i = \langle \dot{n}_i \rangle$ is the mean firing rate of neuron $i$, so that the expansion below is around the first moment of the spike train process. Eq. A5 can be split into free and interacting actions. We expand the action in powers of $\delta \dot{n}_i(t)$ and $\tilde{n}_i(t)$, keeping only terms to quadratic order, which amounts to the Gaussian process approximation,

$$S[\tilde{n}, \delta\dot{n}] \approx \sum_{ij} \int dt dt' \left\{ \tilde{n}_i (\Delta^{-1})_{ij}(t,t') \delta\dot{n}_j - \frac{1}{2}\tilde{n}_i(t)^2 \phi_i \right\}, \tag{A7}$$

where

$$(\Delta^{-1})_{ij}(t,t') \equiv \delta_{ij}\delta(t-t') - \phi_i^{(1)} J_{ij}(t-t') \tag{A8}$$

is the inverse of the linear response function and $\phi_i^{(n)} = \frac{d^n\phi(x)}{dx^n}|_{x=\mu_i+\sum_j J_{ij}r_j}$ is the $n^{\text{th}}$ derivative of the nonlinear activation function evaluated at the mean firing rate $r_i$. The linear order terms have been eliminating by imposing that $r_i$ satisfies

$$r_i = \phi_i \left( \mu_i + \sum_j J_{ij} r_j \right), \tag{A9}$$

and assuming a time-independent solution.

The quadratic action A7 corresponds to a Gaussian distribution for the fluctuations $\delta\dot{n}_i(t)$, which have zero mean and covariance [12]

$$C_{ij}(t', t'') = \sum_k \int_{-\infty}^{\infty} dt \; \Delta_{ik}(t', t)\Delta_{jk}(t'', t) r_k. \tag{A10}$$

For a Gaussian process the moment generating functional of the spike train can be derived [39], giving

$$Z[\tilde{j}] = \int \mathcal{D}\tilde{n}(t)\mathcal{D}\dot{n}(t) e^{\sum_i \int dt \tilde{j}_i(t)\dot{n}_i(t)} \; e^{-S[\tilde{n},\dot{n}]} \approx \exp\left( \sum_i \int dt \; \tilde{j}_i(t) r_i + \frac{1}{2} \sum_{ij} \int dt' dt'' \tilde{j}_i(t') C_{ij}(t', t'') \tilde{j}_j(t'') \right). \tag{A11}$$

Combined with Eq. 5 and Eq. 6, the derived moment generating functional can be used to solve the MLE equations in Eq. 3 and Eq. 4, leading to the closed maximum likelihood estimation equations,

$$C_{ij}(t-t') = r_i \sum_k \int dt'' C_{jk}(t'-t'') \hat{J}_{ik}(t-t''),$$

(Eq. 7 in the main text), which establishes the relationship between the spike train correlation function $C_{ij}$ and the MLE inferred filters $\hat{J}_{ij}$ under Gaussian process approximation.

#### 4. General solution of the integral equation

Eq. 7 is an integral equation for the unknown filters $\hat{J}_{i\ell}(t')$. One might hope to be able to extend the limits of integration to the entire real line and take a Fourier transform to obtain a matrix system of equations that can be solved, but without explicitly imposing the causality constraint this procedure will generally yield a non-causal solution. To use the Fourier method, one first needs to generalize the equation to

$$G_{ij}(t) = r_i \sum_{\ell=1}^{N_{\text{obs}}} \int_{-\infty}^{\infty} dt'' \; \hat{J}_{i\ell}(t'') C_{j\ell}(t''-t), \tag{A12}$$

where

$$G_{ij}(t) = \begin{cases} C_{ij}(t), & t > 0 \\ G^{-}_{ij}(t), & t \leq 0 \end{cases} \tag{A13}$$

for some unknown functions $G^-_{ij}(t)$ that must be determined as part of our solution. Although we have introduced an extra set of unknowns, once we solve for $\hat{J}_{i\ell}(t'')$ the $G^-_{ij}(t)$'s will be determined. This extra set of functions enables causal solutions for the filter by absorbing any non-causal pieces into them. We apply the Fourier transform to obtain

$$G^+_{ij}(\omega) + G^-_{ij}(\omega) = \sum_{\ell=1}^{N_{\rm obs}} r_i \hat{J}^+_{i\ell}(\omega) C_{\ell j}(\omega),$$

where we use $C_{j\ell}(t) = C_{\ell j}(-t)$ and defined the transforms

$$f^+(\omega) = \int_{0+}^{\infty} dt\ e^{-i\omega t} f(t),$$
$$f^-(\omega) = \int_{-\infty}^{0^+} dt\ e^{-i\omega t} f(t),$$
$$f(\omega) = \int_{-\infty}^{\infty} dt\ e^{-i\omega t} f(t).$$

We can write the equation to solve in matrix form,

$$\mathbf{G}^+(\omega) + \mathbf{G}^-(\omega) = \hat{\mathbf{J}}^+(\omega)\mathbf{C}(\omega).$$

Next, we assume we can decompose $\mathbf{C}(\omega) = \mathbf{S}_+(\omega)\mathbf{S}_-(\omega)$, where $\mathbf{S}_+(\omega)$ is analytic and non-vanishing in the upper half plane and $\mathbf{S}_-(\omega)$ is analytic and non-vanishing in the lower half plane.

Continuing, we assume $\mathbf{S}_-(\omega)$ has an inverse, such that we may write

$$\mathbf{G}^+(\omega) \left[\mathbf{S}_-(\omega)\right]^{-1} + \mathbf{G}^-(\omega) \left[\mathbf{S}_-(\omega)\right]^{-1} = \hat{\mathbf{J}}^+(\omega)\mathbf{S}_+(\omega).$$

Next, we split $\mathbf{G}^+(\omega) \left[\mathbf{S}_-(\omega)\right]^{-1} = (\mathcal{F}^{-1}[\mathbf{G}^+ \left[\mathbf{S}_-\right]^{-1})^+(\omega) + (\mathcal{F}^{-1}[\mathbf{G}^+ \left[\mathbf{S}_-\right]^{-1}])^-(\omega)$, where the two terms are defined by first taking the inverse Fourier transform of the left-hand side and then splitting the Fourier transform up into the $\pm$ components. We can then rearrange our equation as

$$(\mathcal{F}^{-1}[\mathbf{G}^+ \left[\mathbf{S}_-\right]^{-1}])^-(\omega) + \mathbf{G}^-(\omega) \left[\mathbf{S}_-(\omega)\right]^{-1} = \hat{\mathbf{J}}^+(\omega)\mathbf{S}_+(\omega) - (\mathcal{F}^{-1}[\mathbf{G}^+ \left[\mathbf{S}_-\right]^{-1})^+(\omega),$$

where by construction the left-hand-side has all of its poles in the lower half plane and the right hand side has all of its poles in the upper half plane. Because the two sides are analytic on different half-planes, the only possibility is that they are both equal to the same function, which must be polynomial of degree $n$ if we require the growth at $|\omega| \to \infty$ to be less than $\mathcal{O}(\omega^n)$ [20]. If we demand that the filters decay as $|\omega| \to \infty$ (which excludes a $\delta$-function component), then the only option is that the two sides must be equal to zero, and hence we arrive at the formal solution

$$\hat{\mathbf{J}}^+(\omega) = (\mathcal{F}^{-1}[\mathbf{G}^+ \left[\mathbf{S}_-\right]^{-1})^+(\omega)[\mathbf{S}_+(\omega)]^{-1}. \tag{A14}$$

In practice, the primary obstacles in performing this procedure are finding a spectral decomposition of the kernel $\mathbf{C}(\omega)$ and then splitting up $\mathbf{G}^+(\omega)[\mathbf{S}_-(\omega)]^{-1}$ into its separate additive factors that are analytic on different half planes. For a one-dimensional system there is a general procedure for performing both of these steps, but for a system of equations the non-commutativity of matrices prohibits the use of the scalar method.

To this end, in this work we will focus our analytic investigations on the case of single neurons or homogeneous networks in order to glean at least some analytic insights into the maximum likelihood inference procedure. This is best illustrated with some concrete examples, which we work through in the next section.

#### 5. Example case: all-to-all coupled network driven by independent noise

To evaluate an explicit example, we consider an all-to-all coupled network with $J_{ij}(t-t') = Jh(t-t')$, for some temporal profile $h(t)$. We evaluate the solutions explicitly for an exponential filter, which has simpler analytic expressions that the alpha functions we use in our simulations, and then give the corresponding results for alpha function filters.

For this all-to-all network the mean field equation reduces to a single rate equation (due to homogeneity of the network),

$$r = \lambda_0 \exp\left(\mu + NJr\right),$$

where $\sum_j \int dt' \; J_{ij}(t-t')r_j$ integrates to $NJr$ for constant $r$. By manipulating this into the form of the Lambert transcendental equation, $W(z)e^{W(z)} = z$ we can write the solution in terms of the Lambert W function,

$$r = -\frac{W_{-1}\left(NJ\lambda_0 e^{\mu}\right)}{NJ}.$$

Next, we can calculate the linear response function by inverting Eq. (A8). This is easiest to do by first Fourier transforming the equation to turn the convolutions in time into multiplications in frequency space, and then solving the matrix equation

$$\sum_{k=1}^{N} \left[\delta_{ik} - gJh(\omega)\right] = \delta_{ij},$$

where $g \equiv \phi'(\mu + NJr)$ is the gain of the network in steady-state (equal to the firing rate $r$ when $\phi$ is exponential), $h(\omega)$ is the Fourier transform of $h(t)$. If we denote $\mathbb{I}$ as the identity and $\mathbf{P}$ as a matrix of all 1's, then the inverse is [41]

$$\left[a\mathbb{I} + b\mathbf{P}\right]^{-1} = \frac{1}{a}\mathbb{I} - \frac{b}{a(Nb+a)}\mathbf{P}.$$

In our case we have $a = 1$, $b = -gJh(\omega)$, giving

$$\Delta_{ij}(\omega) = \delta_{ij} + \frac{gJh(\omega)}{1 - NJgh(\omega)}.$$

Let's now assume an exponential filter $h(t) = \exp(-t/\tau)\Theta(t)/\tau$, which has Fourier transform $h(\omega) = 1/(1+i\omega\tau)$ using the convention $h(\omega) = \int_{-\infty}^{\infty} dt e^{-i\omega t} h(t)$. Thus, in the time-domain $\Delta_{ij}(t)$ is given by

$$\begin{aligned}
\Delta_{ij}(t) &= \int_{-\infty}^{\infty} \frac{d\omega}{2\pi} e^{i\omega t}\left(\delta_{ij} + \frac{J}{h(\omega)^{-1} - NJg}\right) \\
&= \int_{-\infty}^{\infty} \frac{d\omega}{2\pi} e^{i\omega t}\left(\delta_{ij} + \frac{gJ}{i\omega\tau + 1 - NJg}\right) \\
&= \delta_{ij}\delta(t) + gJ\exp(-(1-NJg)t/\tau)\Theta(t)/\tau,
\end{aligned}$$

where to evaluate the second term we used the residue theorem: factoring out a $i\tau$ from the denominator, we observe a pole at $\omega = i(1-NJg)$ in the upper-half plane when $1 > NJg$. This restriction requires either $0 < Jg < 1/N$ or $J < 0$ for the process to be stable. For $t < 0$, $i\omega t = -iR|t|(\cos\theta + i\sin\theta)$ on a contour of radius $R$, and the real part of this, $+R|t|\sin\theta$ is only negative in the lower-half plane, so we must close the contour there and the integral evaluates to zero because there are no poles contained in the contour. For $t > 0$ the real part of the arc is $-R|t|\sin\theta$, and we must close the arc in the upper half plane, obtaining the contribution from the pole. Note that the response function is causal in time.

With the response function in hand we may use Eq. A10 to evaluate the covariance for this model. In the Fourier domain we have

$$
\begin{aligned}
C_{ij}(\omega) &= \sum_{k=1}^{N} \Delta_{ik}(\omega)\Delta_{jk}(-\omega) r \\
&= r\sum_{k=1}^{N}\left(\delta_{ik} + \frac{J}{i\omega\tau + 1 - NJg}\right)\left(\delta_{jk} + \frac{Jg}{-i\omega\tau + 1 - NJg}\right) \\
&= r\sum_{k=1}^{N}\left(\delta_{ik}\delta_{jk} + \frac{J\delta_{jk}}{i\omega\tau + 1 - NJg} + \frac{Jg\delta_{ik}}{-i\omega\tau + 1 - NJg} + \frac{J^2g^2}{|i\omega\tau + 1 - NJg|^2}\right) \\
&= r\left[\delta_{ij} + 2Jg\mathrm{Re}\left[\frac{1}{i\omega\tau + 1 - NJg}\right] + \frac{NJ^2}{(1-NJg)^2 + (\omega\tau)^2}\right] \\
&= r\left[\delta_{ij} + \frac{2Jg(1-NJg) + NJ^2g^2}{(i\omega\tau + 1 - NJg)(-i\omega\tau + 1 - NJg)}\right] \\
&= r\left[\delta_{ij} + \frac{Jg(2-NJg)}{(i\omega\tau + 1 - NJg)(-i\omega\tau + 1 - NJg)}\right]
\end{aligned}
$$

We again evaluate the inverse Fourier transform by using the Residue theorem. There are now two symmetric poles at $\omega = \pm i(1-NJg)/\tau$, so we get a contribution from both planes, as expected for a covariance. The result is

$$
C_{ij}(t) = r\left[\delta_{ij}\delta(t) + \frac{Jg(2-NJg)}{2(1-NJg)}\frac{\exp(-(1-NJg)|t|/\tau)}{\tau}\right].
$$

We can use this result with Eq. 7 to solve for the inferred filters $\hat{J}(t)$.

### 6. Solution for a single observed unit

Eq. 7 is a Wiener-Hopf integral equation that is difficult to solve in the multivariate case, but is tractable in the scalar case, which corresponds to a single observed neuron in our context. We calculate this here for unit $i = 1$ in the all-to-all network. The equation to solve is

$$
G^{+}(\omega) + G^{-}(\omega) = r\hat{J}^{+}(\omega)C(\omega),
$$

where

$$
\begin{aligned}
G^{+}(\omega) &= r\int_{0^+}^{\infty} d\tau\ e^{-i\omega\tau}\left(\delta(\tau) + \frac{a}{2b\tau}e^{-b|t|/\tau}\right) \\
&= \frac{ar}{2b}\frac{1}{i\omega\tau + b},
\end{aligned}
$$

where we introduce $a = Jg(2-NJg)$ and $b = 1-NJg$ to simplify the upcoming formulas. The full Fourier transform of $C(\omega)$

$$
C(\omega) = r\left[1 + \frac{a}{(i\omega\tau + b)(-i\omega\tau + b)}\right].
$$

Therefore, we need to solve the equation

$$\begin{aligned}\frac{a}{2b}\frac{1}{i\omega\tau+b}+r^{-1}G^{-}(\omega)&=r\hat{J}^{+}(\omega)\left(1+\frac{a}{(i\omega\tau+b)(-i\omega\tau+b)}\right)\\&=r\hat{J}^{+}(\omega)\left(\frac{(i\omega\tau+b)(-i\omega\tau+b)+a}{(i\omega\tau+b)(-i\omega\tau+b)}\right)\\&=r\hat{J}^{+}(\omega)\left(\frac{-(i\omega\tau)^2+b^2+a}{(i\omega\tau+b)(-i\omega\tau+b)}\right)\\&=r\hat{J}^{+}(\omega)\left(\frac{(i\omega\tau+\sqrt{b^2+a})(-i\omega\tau+\sqrt{b^2+a})}{(i\omega\tau+b)(-i\omega\tau+b)}\right),\end{aligned}$$

where we divided both sides by one factor of $r$. We now separate the factors that are analytic and non-vanishing on the lower-half-plane and the upper half-planes. We have

$$\frac{a}{2b}\frac{1}{i\omega\tau+b}\frac{-i\omega\tau+b}{-i\omega\tau+\sqrt{b^2+a}}+r^{-1}G^{-}(\omega)\frac{-i\omega\tau+b}{-i\omega\tau+\sqrt{b^2+a}}=r\hat{J}^{+}(\omega)\left(\frac{i\omega\tau+\sqrt{b^2+a}}{i\omega\tau+b}\right).$$

We use partial fractions on the left-hand-side to write

$$\frac{a}{2b}\frac{1}{i\omega\tau+b}\frac{-i\omega\tau+b}{-i\omega\tau+\sqrt{b^2+a}}=\frac{A}{i\omega\tau+b}+\frac{B}{-i\omega\tau+\sqrt{b^2+a}},$$

where

$$A=\frac{a}{b+\sqrt{b^2+a}},\ B=-\frac{a}{2b}\frac{\sqrt{b^2+a}-b}{\sqrt{b^2+a}+b}.$$

Here we will only care about the filter $\hat{J}^{+}(\omega)$, so we only need the $A$ term. After separating the terms analytic in the upper versus lower half planes, demanding that the filters decay at infinite $\omega$ means we must have

$$\begin{aligned}r\hat{J}^{+}(\omega)\left(\frac{i\omega\tau+\sqrt{b^2+a}}{i\omega\tau+b}\right)&=\frac{a}{b+\sqrt{b^2+a}}\frac{1}{i\omega\tau+b}\\\Rightarrow r\hat{J}^{+}(\omega)&=\frac{a}{b+\sqrt{b^2+a}}\frac{1}{i\omega\tau+\sqrt{b^2+a}}\end{aligned}$$

Because $\hat{J}^{+}(\omega)$ only has poles in the lower half plane, as desired, we know it will be causal and we can use the regular Fourier transform to recover it in the time domain (as $\hat{J}^{+}(\omega)=\hat{J}(\omega)$). The result is

$$\hat{J}(t)=\frac{1}{r}\frac{a}{b+\sqrt{b^2+a}}\frac{e^{-\sqrt{b^2+a}\ t/\tau}}{\tau}\Theta(t).$$

Restoring $a=Jg(2-NJg)$ and $b=1-NJg$ gives

$$\hat{J}(t)=\frac{g}{r}\frac{J(2-NJg)}{1-NJg+\sqrt{(1-NJg)^2+Jg(2-NJg)}}\frac{e^{-\sqrt{(1-NJg)^2+Jg(2-NJg)}\ t/\tau}}{\tau}\Theta(t).\tag{A15}$$

We can check the limit of the fully resolved case when $N=1$. We have $(1-Jg)^2+Jg(2-Jg)=1-2Jg+(Jg)^2+2Jg-(Jg)^2=1$, giving $\hat{J}(t)=g/rJe^{-t/\tau}\Theta(t)/\tau$. This shows that when the nonlinearity of the generative model is not exponential (a model-mismatch) the inferred filter is off from the ground truth by a multiplicative factor $g/r$. For the model-matched case where both the generative model and the inference model use an exponential nonlinearity the gain is equal to the rate, $g=r$, and we recover the true filter. We can also verify in Mathematica that this solution does satisfy the original integral equation.

Now that we have $\hat{J}(t)$ and $C(t)$ we can evaluate the normalized overlap between them,

$$\rho = \frac{\int_0^\infty dt\ \hat{J}(t)C(t)}{\sqrt{\int_0^\infty dt\ \hat{J}(t)^2 \int_0^\infty dt\ C(t)^2}}.$$

Using Mathematica, this works out to

$$\rho = \frac{2\sqrt{1-NJr}\sqrt[4]{(1-NJr)^2 + Jr(2-NJr)}}{1 - NJr + \sqrt{(1-NJr)^2 + Jr(2-NJr)}}$$

for the model-matched case $g = r$. Plotting this as a function of $x = NJr \in [0, 1)$ for fixed $N$, we see that for small $x$ $\rho \approx 1$, and rapidly approaches 0 as $x \to 1$ from below. As $N$ increases the fraction of the range of $x$ for which $\rho \approx 1$ increases.

#### a. $N_{\text{obs}}$ observed neurons

For the homogeneous all-to-all network we can also exactly solve for the inferred synaptic connections, as all connection filters will be the same, $\hat{J}_{rr'}(t) = \hat{J}(t)$. The equation we must solve becomes

$$\frac{a}{2b}\frac{1}{i\omega\tau + b} + r^{-1}G^-(\omega) = r\hat{J}^+(\omega)\left(1 + \frac{N_{\text{obs}}a}{(i\omega\tau + b)(-i\omega\tau + b)}\right).$$

Following the same steps as above, we obtain the inferred filters

$$\hat{J}(t) = \frac{g}{r}\frac{J(2-NJg)}{1 - NJg + \sqrt{(1-NJg)^2 + N_{\text{obs}}Jg(2-NJg)}}\frac{e^{-\sqrt{(1-NJg)^2 + N_{\text{obs}}Jg(2-NJg)}\ t/\tau}}{\tau}\Theta(t). \tag{A16}$$

The reader can check that we recover the ground truth filter when $N_{\text{obs}} \to N$ and $g \to r$.

#### b. Alpha function filter

The manipulations work similarly for an alpha function filter $h(t) = te^{-t/\tau}\Theta(t)/\tau^2$, which has Fourier transform $h(\omega) = 1/(i\omega\tau + 1)^2$. This introduces more poles to deal with when using the residue theorem and partial fraction decomposition, but the calculations are tractable for the most part. We find the covariance of the units to be

$$C_{ij}(t-t') = r\left[\delta_{ij}\delta(t-t') + \frac{(a_-(1)^2 - b_+^2)(b_+^2 - a_+(1)^2)}{(b_+ - b_-)(b_+ + b_-)}\frac{e^{-b_+|t-t'|/\tau}}{2b_+\tau} - \frac{(a_-(1)^2 - b_-^2)(b_-^2 - a_+(1)^2)}{(b_+ - b_-)(b_+ + b_-)}\frac{e^{-b_-|t-t'|/\tau}}{2b_-\tau}\right], \tag{A17}$$

where $a_\pm(n) = \sqrt{1 + (N-n)Jg \pm \sqrt{(N-n)Jg(4-Jg)}}$ and $b_\pm = 1 \pm \sqrt{NJg}$, and the effective self-history filter to be

$$\hat{J}(t) = \frac{1}{r}\left[A_+(N_{\text{obs}})e^{-a_+(N_{\text{obs}})t/\tau} - A_-(N_{\text{obs}})e^{-a_-(N_{\text{obs}})t/\tau}\right]\frac{\Theta(t)}{\tau}. \tag{A18}$$

The full expressions for the amplitudes are

$$
\begin{aligned}
DA_+(N_{\rm obs}) = {} & b_-^3 b_+^3 + b_-^2 b_+^2 a_-(1)^2 + b_-^2 b_+^2 a_+(1)^2 - b_-^2 a_-(1)^2 a_+(1)^2 - b_- b_+ a_-(1)^2 a_+(1)^2 - b_+^2 a_-(1)^2 a_+(1)^2 \\
& + (b_- + b_+)(b_-^2 b_+^2 - a_-(1)^2 a_+(1)^2) a_+(N_{\rm obs}) \\
& + a_-(N_{\rm obs})^2 \Big\{ - b_-^3 b_+ - b_-^2 b_+^2 + a_-(1)^2 a_+(1)^2 + b_- b_+ (-b_+^2 + a_-(1)^2 + a_+(1)^2) \\
& \qquad - (b_- + b_+)(b_-^2 + b_+^2 - a_-(1)^2 - a_+(1)^2) a_+(N_{\rm obs}) \Big\}, \qquad \text{(A19)} \\
DA_-(N_{\rm obs}) = {} & b_-^3 b_+^3 + b_-^2 b_+^2 a_-(1)^2 + b_-^2 b_+^2 a_+(1)^2 - b_-^2 a_-(1)^2 a_+(1)^2 - b_- b_+ a_-(1)^2 a_+(1)^2 - b_+^2 a_-(1)^2 a_+(1)^2 \\
& + (b_- + b_+)(b_-^2 b_+^2 - a_-(1)^2 a_+(1)^2) a_+(N_{\rm obs}) \\
& + a_-(N_{\rm obs})^2 \Big\{ - b_-^3 b_+ - b_-^2 b_+^2 + a_-(1)^2 a_+(1)^2 + b_- b_+ (-b_+^2 + a_-(1)^2 + a_+(1)^2) \\
& \qquad - (b_- + b_+)(b_-^2 + b_+^2 - a_-(1)^2 - a_+(1)^2) a_+(N_{\rm obs}) \Big\}. \qquad \text{(A20)}
\end{aligned}
$$

where $D = (b_- + a_-(N_{\rm obs}))(b_- + a_+(N_{\rm obs}))(b_+ + a_-(N_{\rm obs}))(b_+ + a_+(N_{\rm obs}))(a_-(N_{\rm obs}) - a_+(N_{\rm obs}))$. As stated in the main text, when $N_{\rm obs} = 1$ these simplify to

$$
\begin{aligned}
A_+(1) &= \frac{(a_+(1) - b_-)(a_+(1) - b_+)}{a_+(1) - a_-(1)}, \\
A_-(1) &= \frac{(a_-(1) - b_-)(a_-(1) - b_+)}{a_+(1) - a_-(1)}.
\end{aligned}
$$
